# Dynamic patterns of transcript abundance of transposable element families in maize

**DOI:** 10.1101/668558

**Authors:** Sarah N Anderson, Michelle C Stitzer, Peng Zhou, Jeffrey Ross-Ibarra, Cory D Hirsch, Nathan M Springer

**Author notes:** **Email addresses:**, (S.N.A.), (M.C.S.);, (P.Z.);, (J.R.I);, (C.D.H);, (N.M.S.). **Corresponding author**: Nathan M. Springer, 140 Gortner Laboratory, 1479 Gortner Ave., St. Paul, MN, 55108.

## Abstract

Transposable Elements (TEs) are mobile elements that contribute the majority of DNA sequences in the maize genome. Due to their repetitive nature, genomic studies of TEs are complicated by the difficulty of properly attributing multi-mapped short reads to specific genomic loci. Here, we utilize a method to attribute RNA-seq reads to TE families rather than particular loci in order to characterize transcript abundance for TE families in the maize genome. We applied this method to assess per-family expression of transposable elements in >800 published RNA-seq libraries representing a range of maize development, genotypes, and hybrids. While a relatively small proportion of TE families are transcribed, expression is highly dynamic with most families exhibiting tissue-specific expression. A large number of TE families were specifically detected in pollen and endosperm, consistent with reproductive dynamics that maintain silencing of TEs in the germ line. We find that B73 transcript abundance is a poor predictor of TE expression in other genotypes and that transcript levels can differ even for shared TEs. Finally, by assessing recombinant inbred line and hybrid transcriptomes, complex patterns of TE transcript abundance across genotypes emerged. Taken together, this study reveals a dynamic contribution of TEs to maize transcriptomes.

## Introduction

Plant genomes contain an abundance of transposable elements (TEs) which can increase in copy number through transposition within a host genome. TEs are broadly classified into two classes based on whether they utilize a DNA or RNA intermediate for movement. Classes are further divided into orders based on transposition mechanisms and then into superfamilies based on structural features. Within each superfamily, TE family classifications are based on sequence similarity, particularly at the terminal repeats (Wicker *et al.* 2007; Stitzer *et al.* 2019). Individual TEs can be described as autonomous if they code for all enzymes required for transposition or non-autonomous if one or more of these sequences is missing. Since TE proteins can act in trans, autonomous members of a family can allow for transposition of other autonomous or non-autonomous elements of the same family. The sequence similarity of families along with family-dependent variability in distinct genomic distributions (Stitzer *et al.* 2019) and methylation patterns (Eichten *et al.* 2012) make TE family the preferred level for analysis of groups of TEs on the genomic scale.

Due to the potential detrimental consequences of unchecked transposition, the host genome has employed mechanisms to constrain TE movement. The analysis of chromatin modifications suggests that most TEs are associated with heterochromatin modifications and thus transcriptionally suppressed (Rabinowicz *et al.* 1999; Yuan *et al.* 2002; West *et al.* 2014). Additionally, silencing at some loci is reinforced through RNA-directed DNA methylation (RdDM) where transcripts are processed into small RNAs that can act in trans to direct DNA methylation. Therefore, transcription of TEs is associated with both active TEs, which require full transcripts to facilitate movement, and actively silenced TEs, where even partial transcripts can trigger small RNA production.

The most conclusive evidence of functional transcription of TEs is actually indirect -- the generation of novel TE insertions requires transcription and translation of a TE encoded protein. Classical genetic studies have found evidence for active TE families in some maize germplasm (Robertson 1978; McClintock 1950). These DNA terminal inverted repeat (TIR) transposons require expression of a functional transposase from an autonomous element for the transposition of both autonomous and non-autonomous elements. There is evidence that the expression of these transposase genes can be influenced by copy number (Fußwinkel *et al.* 1991; Rudenko and Walbot 2001) as well as epigenetic regulation (Lisch and Bennetzen 2011). There is also evidence that stress, such as tissue culture, can result in activation of expression and subsequent transposition of DNA transposons in maize and other species (Peschke *et al.* 1987). LTR retrotransposons, however, require expression of the terminal repeats plus internal (oftentimes protein coding) domains for transposition. While the maize genome has a large number of LTRs including many young elements (Stitzer *et al.* 2019) there have been few examples of new mutations resulting from LTR elements (Wessler and Varagona 1985; Jin and Bennetzen 1989; Varagona *et al.* 1992). Detection of novel insertions of active LTR transposons has been limited in maize, with only a small number of recent transposition events detected in large genetic screens (Varagona *et al.* 1992; Dooner *et al.* 2019). There is evidence for reactivation of retrotransposons by tissue culture in tobacco and rice (Grandbastien *et al.* 1989; Pouteau *et al.* 1991; Hirochika *et al.* 1996), and transcripts for some of these families can also be induced through other environmental stresses (Grandbastien 2004). Further, in *Arabidopsis* there is evidence for activation of some TEs in the vegetative nucleus of male gametophytes (Slotkin *et al.* 2009). Expression, and in some cases movement, of DNA transposons and retrotransposons can also be the result of mutations in genes that regulate DNA methylation or other chromatin modifications (Miura *et al.* 2001; Kato *et al.* 2003; Woodhouse *et al.* 2006; Reinders *et al.* 2009; Mirouze *et al.* 2009; Anderson *et al.* 2018).

While TE expression in maize can be inferred from active transposition and analysis of individual TE transcripts, assessment of transcript abundance of TEs on a genomic scale has been limited. Analysis of EST sequences with homology to TEs revealed dynamic expression of some TE families in maize tissues (Vicient 2010), and assessment of RNA-seq data has revealed some expression variation among different TE types in the maize genome (Diez *et al.* 2014). However, it remains challenging to utilize short-reads derived from RNA-seq experiments to assess transcript abundance of TEs due to difficulties analyzing repetitive sequences that do not map uniquely to the genome (Slotkin 2018), and interpretations of results can differ based on methods used (Bousios *et al.* 2017). Despite these challenges, assessing TE transcript abundance has the potential to reveal substantial insights into how variable TEs can influence host genomes. Since TEs can contain regulatory elements capable of influencing both the TE itself and neighboring gene expression, expressed TEs may denote candidates for functional relevance to gene regulation (Makarevitch *et al.* 2015; Oka *et al.* 2017; Zhao *et al.* 2018). In this study, we describe and implement an approach that allows for analysis of the expression of TE families through mapping to the complex genome of maize. By monitoring per-family expression levels we can survey existing RNA-seq data to determine broad properties of transcript accumulation of maize TEs. This revealed that while only a small proportion of all TE families are expressed, TE expression is dynamic across development, genotypes, and hybrids.

## Materials and Methods

### Data sources

RNA-seq data for all samples were obtained from published datasets (Zhou *et al.* 2019; Li *et al.* 2012, 2013; Stelpflug *et al.* 2016; Walley *et al.* 2016; Lin *et al.* 2017). Libraries were downloaded from SRA, and a table of libraries and accession numbers can be found in Table S1.

### TE expression analysis

RNA-seq libraries were processed by trimming with cutadapt v.1.8.1 (-m 30 -q 10 --quality-base=33) following by mapping to the B73v4 genome assembly (Jiao *et al.* 2017) with tophat2 v.2.0.13 (-g 20 -i 5 -I 60000) (Kim *et al.* 2013), allowing for up to 20 mapping positions. BAM output files were then sorted, converted to SAM format, and reformatted for compatibility with HTSeq (Anders *et al.* 2015) using the convert_sam_to_all_NH1_v2.pl script. Using HTseq v.0.5.3, reads were intersected with a modified annotation file B73.structuralTEv2.1.07.2019.filteredTE.subtractexon.plusgenes.chr.sort.gff3. This annotation file was created by first masking exons from the disjoined TE annotation file B73.structuralTEv2.2018.12.20.filteredTE.disjoined.gff3 found at https://github.com/SNAnderson/maizeTE_variation, appending full gene model annotations, and reformatting to remove features on contigs and to format for use in HTseq. TE and gene expression was counted from the SAM output of HTseq using script te_family_mapping_ver6.pl.

Briefly, reads were assigned to genes when they map uniquely and hit a gene annotation but not any TE annotation. Reads were assigned to a TE element when they map uniquely and hit a single TE annotation (with an overlap of at least 1 bp), and reads were assigned to a TE family when they map uniquely or multiple times but only hit a single TE annotation. Ambiguous reads were defined by hits assigned to both a gene and a TE and were counted to columns labeled te.g for the TE element or family. Two output files were created for each library. The first file contains TE family counts for 4 categories of reads: unique reads hitting one TE family (u_te.fam; Read E in Figure 1), unique ambiguous reads (u_te.g; Read B in Figure 1), multi reads to one family (m_te.fam; Read D in Figure 1), and multi ambiguous reads (m_te.g; Read C in Figure 1). The second file contains unique counts only to both genes and TEs and contains two columns: unique reads hitting a single TE or gene (unique; Reads A and E in Figure 1) and unique reads hitting both a TE and a gene (te.g; Read B in Figure 1). In addition, a single line for each library with the total number of reads assigned to each category is added to a file called te_mapping_summary.txt.

**Figure 1.**
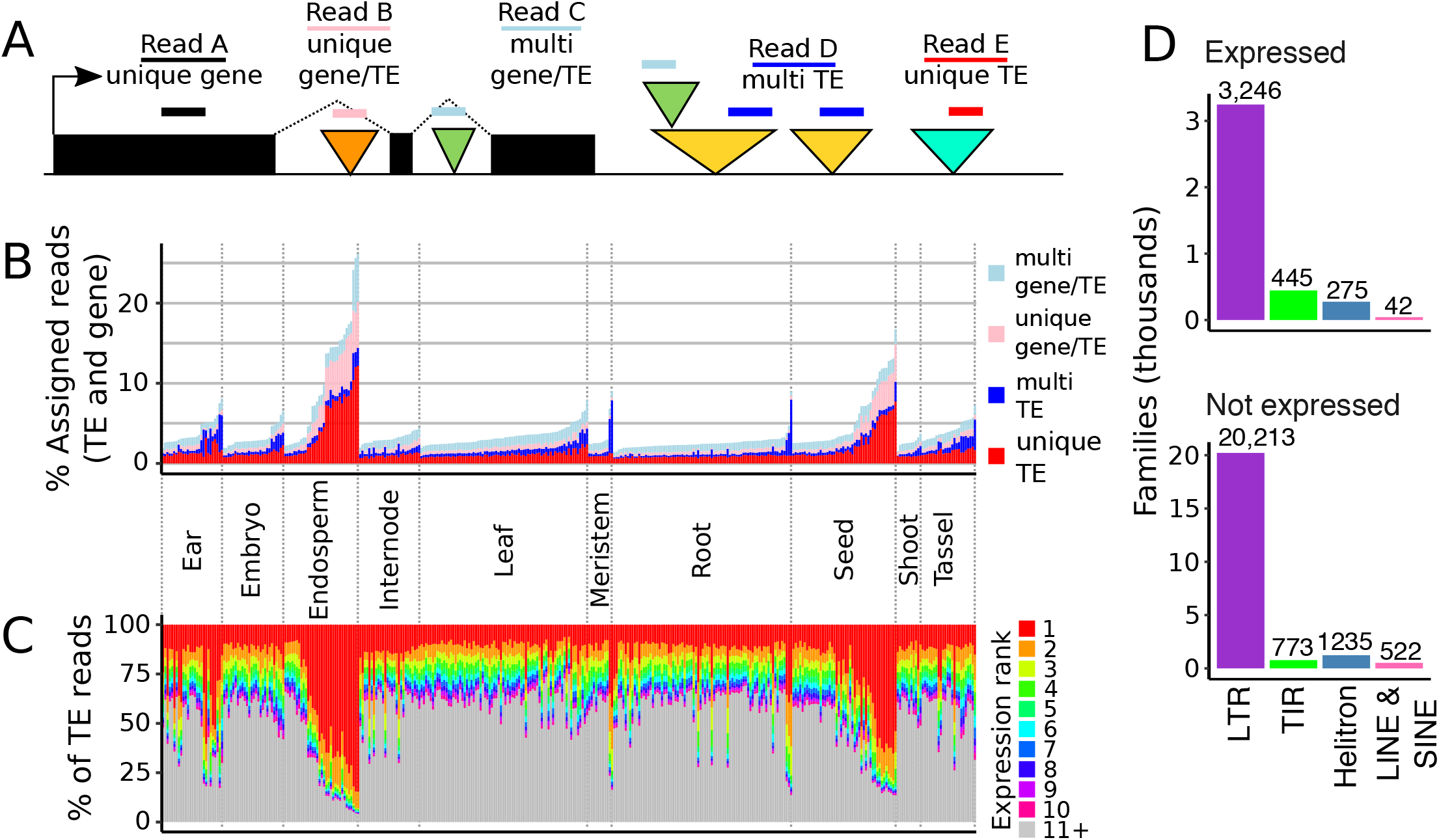

Count tables for each library were then combined using the script combine_count_totals.pl. Four output tables were created. The first file (multi_combined_counts.txt) includes all four count columns per library for each TE family and the second file (element_combined_counts.txt) includes two count columns per library for each TE element and gene. The third file (family_sum_combined_counts.txt) contains counts for TE families in a single column per library, corresponding to the sum of u_te.fam and m_te.fam columns. Finally, the fourth file (family_prop_unique.txt) contains a single column per library with the proportion of the reads in file three that are derived from unique-mapping reads.

Unless otherwise noted, all scripts and files referred to can be found at https://github.com/SNAnderson/maizeTEexpression. See sample_shell_script.sh for full workflow example.

### Expression normalization and differential expression

Expression for genes and TE families was normalized by calculating reads per million (RPM) using the total number of reads assigned to TE families or genes as the denominator. Expression of individual elements was normalized as RPM using the same library size estimate. In the analysis of all B73 expression, genes and TE families were considered expressed where RPM values were > 1 in at least 3 libraries. Only expressed families were used in PCA and tau analyses. PCA was performed using the prcomp function in R and tissue-specificity was estimated with the tau metric using log2(1 + RPM) transformed expression values for genes and TE families. Per-family expression dynamics were visualized in R using pheatmap 1.0.10, with relative expression calculated by dividing transformed expression values by the maximum value for each row. Tau values for expressed genes and TE families were calculated from transformed expression values in R using the fTau function published in (Kryuchkova-Mostacci and Robinson-Rechavi 2017). For the subset of tissues, families were considered expressed if the mean across replicates was at least 1 RPM, and mean values were also used to identify families expressed across the subset or in only one tissue type. Expressed families in the NAM lines were defined with an RPM cutoff of 1 in tissues with no biological replicates, or a mean value of 1 in meristem, which had two biological replicates.

Differential expression (DE) analysis was performed using the R package DESeq2 (Love *et al.* 2014), with normalization performed using the estimateSizeFactors function and adjusted p-values calculated with the FDR method. DE TE families were defined using an FDR cutoff of < 0.05 and a fold-change cutoff of 2. Non-additive expression was defined by significantly higher or lower expression in the F1 than in both parents.

### Unimodal expression in RILs

To test for the segregation of expression values in RILs, Hardigan’s dip test index (Hartigan and Hartigan 1985) was calculated for each DE TE family using the diptest package in R. P-values were calculated using the simulate.p.value option, which computes p-values by Monte Carlo simulation. The null hypothesis of the test is that the distribution of values is non-unimodal (at least bimodal), so families were considered unimodal when the p-value was < 0.05.

### Data availability

All data used in this study are previously published and available in SRA and a table of accession numbers for each library has been uploaded onto figshare. Scripts for performing the TE expression method are available at https://github.com/SNAnderson/maizeTEexpression.

## Results

### TE family analysis captures expression of repetitive sequences

TE sequences are highly repetitive in the maize genome. Due to short read length a considerable fraction of reads from RNA sequencing experiments can not be uniquely mapped to the reference genome. However, since TE families are connected by lineage and sequence similarity, we developed a method to assess per-family TE expression (Figure 1A). Briefly, reads were mapped to the genome using Tophat2 (Kim *et al.* 2013) with the -g 20 option, which reports up to 20 mapping locations for each read. Uniquely mapping reads (such as Read A) that aligned to annotated genes were used to document gene expression levels. Per-family transcript abundance for TEs was determined using both reads that mapped uniquely to a specific TE (read E) as well as reads that mapped to multiple locations that are all annotated as members of the same TE family (read D). In some other cases a uniquely mapping read (read B) or a multiple-mapping read (read C) aligned to a TE that is located within a gene, resulting in ambiguous assignments that can not be fully clarified as gene- or TE-derived. The reads with ambiguous gene/TE assignments (Reads B and C) were summarized per library but removed from downstream analyses. Transcript abundance was normalized as reads per million (RPM), with the total library size determined from the sum of assigned reads to either genes or TEs. The analyses in this manuscript utilized existing RNAseq datasets that focused on polyadenylated transcripts. We will refer to the observed abundance of transcripts for TE families as TE ‘expression’, but it is important to caution that this includes a mix of multiple processes, including transcription of functional TE products, read-through transcription from nearby genes, and non-coding transcripts derived from cryptic promoters within a TE. Importantly, this means that the presence of transcripts does not necessarily imply production of functional products or potential movement of a TE family.

We used this method to assess transcript abundance for TE families in 359 RNAseq libraries from 3 published datasets of B73 inbred plants representing a diverse set of tissues and developmental stages (Zhou *et al.* 2019; Stelpflug *et al.* 2016; Walley *et al.* 2016). TEs accounted for 1.4 to 26.1% of the reads assigned to genes or TEs, with particularly high TE expression seen in later stages of endosperm development (Figure 1B). To distinguish between global up-regulation of many TE families or up-regulation of a small number of TE families we assessed the transcript abundance for the top expressed TE families in each tissue (Figure 1C). This revealed a strong positive correlation between the total TE expression in a library and the expression of the top most highly expressed family in that library (pearson’s correlation = .853, p value < 0.001) suggesting that higher levels of expression in some tissues is due to the increased abundance of one, or several, TE families rather than up-regulation of many families. The largest contribution of a single family was observed in late endosperm and seed tissue, where over half of TE transcripts were assigned to LTR retrotransposon family RLC11137.

Transcripts were detected for 4,008 of the 26,751 TE families in maize, including 3,246 LTR, 445 TIR, 275 Helitron, 27 LINE, and 15 SINE families (Figure 1D). TE families in maize range in size from a single member (20,256 families) to over 16,000 members for the LTR family cinful-zeon (RLG00001). Across all TE orders, larger families often have expression above the 1 RPM threshold. All 30 families with at least 1,800 members and 89% of large families (>500 members) have detectable expression, while only 12% of single member families were expressed (Figure S1). However, since values are not normalized for the length of the elements or number of copies, high copy families with lowly detectable transcripts could represent transcriptional noise. Due to the large number of single-member TE families in the maize genome, the majority of all expressed families have a single member. The TE families that have polyadenylated transcripts do not show strong enrichments or depletions for coding potential or relative age.

### TE expression is highly dynamic across development

Gene expression is known to be highly dynamic in different developmental stages or tissues. PCA plots were created using expression values from genes or TE families to assess the ability of TE family transcript abundance information to capture differences between libraries (Figure S2). Both gene and TE family transcript abundance values clustered by tissue type, however both PC1 and PC2 represented more of the variation between libraries for genes than TE families. To quantify the dynamics of expression across libraries, the tau metric was calculated for each gene and TE family (Yanai *et al.* 2005; Kryuchkova-Mostacci and Robinson-Rechavi 2017) as in (Stitzer *et al.* 2019). Tau values range from 0 to 1, with low numbers representing constitutive expression and high numbers indicating tissue-specific expression. While genes have a bimodal distribution with a similar number of genes showing high and low tau values, TE families are largely skewed to higher tau values (Figure 2A), indicating greater tissue-specificity for the transcript abundance of TEs than for genes. Among TE orders, the distribution of tau values is lower for TIR families than LTR and Helitron families. Interestingly, the tau distributions for large TE families of all orders skews lower than small and single-copy families. This could reflect low levels of transcription for some members of these large families or could reflect different members with different patterns of expression that result in an apparent constitutive expression for the family.

**Figure 2.**
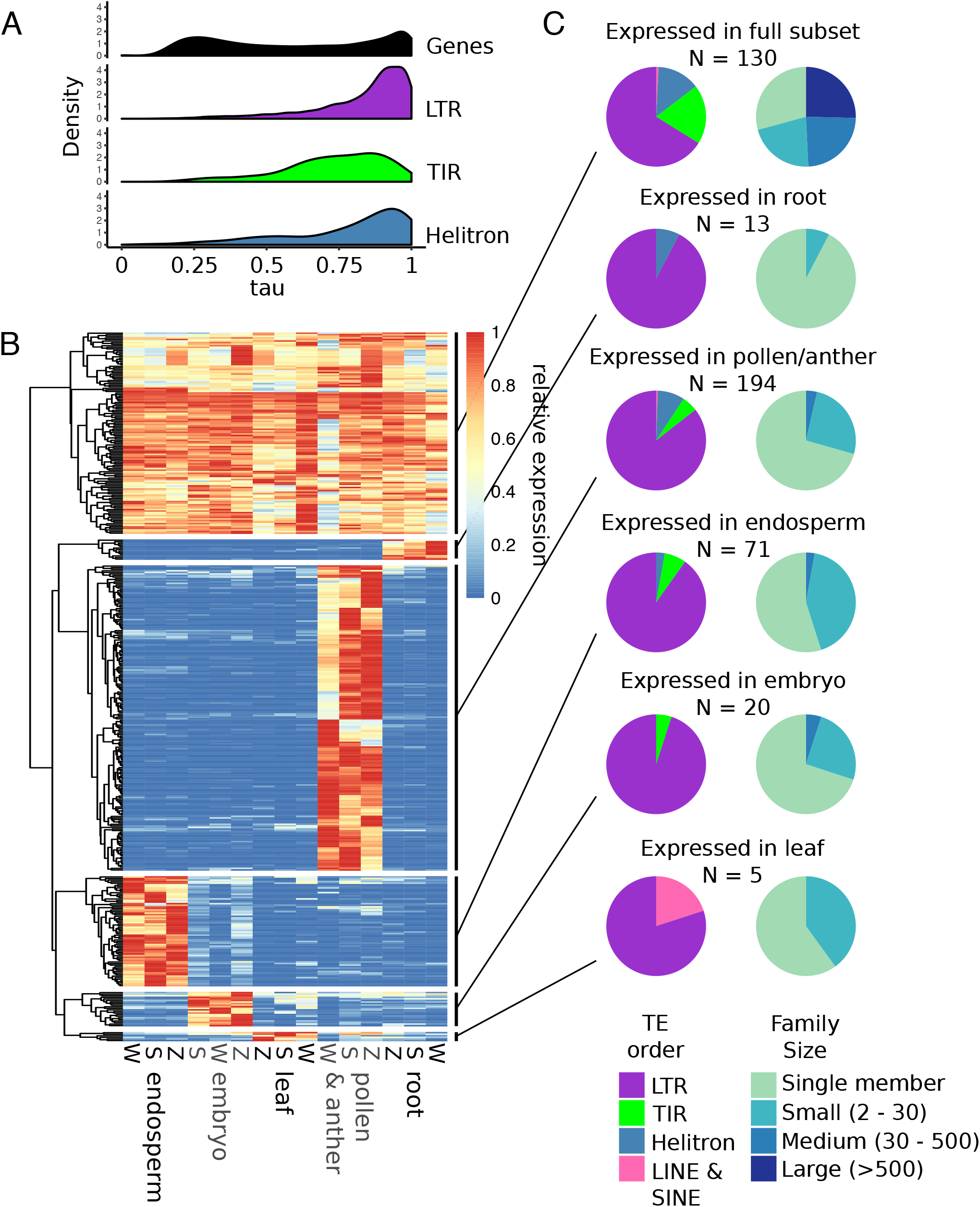

To further assess TE expression dynamics across tissues, a subset of libraries were selected to represent a range of vegetative and reproductive tissues sampled similarly in all three developmental atlases. The tissues selected included leaf, root, embryo, early endosperm, and pollen/anther/floret. The average transcript abundance value for biological replicates was calculated, and families with an average of at least 1 RPM in at least one sample were considered expressed. This identified 2,735 expressed families. A heatmap of relative expression across samples shows that while some families are expressed in most samples, a large number of families are expressed in a single sample or tissue type, particularly in pollen/anther (Figure S3A). TE families expressed across all libraries and those expressed specifically in a single tissue type were selected for further analysis (Figure 2B). There were 130 TE families expressed across the subset of tissues, including LTR, TIR, and Helitron families. An additional 303 families were found to be expressed in a single tissue type. The largest number of tissue-specific families were found in pollen (194 families) or endosperm (71 families), suggesting that de-repression of TEs in these tissue types is specific to unique TE families rather than a global change in TE expression. A smaller number of TE families were specifically expressed in embryo, leaf, or root (20, 5, and 13 families, respectively). Across tissues, LTR retrotransposons represented the largest proportion of expressed families. Tissue-specific families are predominantly small (< 30 members), while nearly half of families expressed across tissues have at least 30 members (Figure 2).

We assessed the relative contribution of unique and multi-mapping reads to the transcript abundance of TE families within this subset of tissues. For each family with more than one member, the proportion of assigned reads that mapped uniquely was averaged across expressed libraries, revealing that, for the majority of families (629 of 1176) unique mapping reads contribute >90% of reads (Figure S3B). However, there were also 102 families where < 20% of reads were uniquely mapped, including 78 LTR, 17 TIR, 6 Helitron, and 1 SINE family. For families expressed across the subset and where the majority of reads could be uniquely mapped to a single element, per-element dynamics of expression could be assessed. This revealed several patterns of transcript abundance, exemplified by three examples in Figure S3. For some families, unique mapped reads suggest transcripts predominantly from a single member (Figure S3C), while other families show transcripts from multiple members at similar frequencies across tissues (Figure S3D). In other cases, expressed elements varied across tissues (Figure S3E).

### TE expression dynamics across genotypes

Transcripts were only observed for a small proportion of TE families present in B73. To assess potential genetic variation for transcription of TE families we assessed TE expression within five tissues of the 26 maize genotypes used as founders in the Nested Association Mapping (NAM) population (Li *et al.* 2012; Lin *et al.* 2017). Transcripts were detected for approximately 1,000 TE families in each library, with little variation in the number of expressed TE families across genotypes despite mapping of all reads to the B73 reference genome (Figure 3A; S4). For each expressed TE family, the number of genotypes with transcripts was assessed revealing that, while families expressed in B73 tend to be commonly expressed in at least 20 genotypes, there are also a large number of TE families with rare expression in < 5 genotypes (Figure 3B). Across all genotypes, nearly 4,000 TE families are expressed in immature ear tissue, with the majority of expressed families exhibiting rare expression (Figure 3C). This pattern holds true across all 5 tissues with expression data for these genotypes (Figure S4). Different maize genotypes have highly variable TE insertions (Anderson *et al.* 2019), so variation in expressed TE families may result from differences in the number of TE family members among genomes, variation in chromatin state around particular TE insertions, or differential abundance of trans-acting factors. The observation of expression of TE families in other genotypes does not necessarily imply activation of an element that is present in B73, but may instead reflect expression of a novel member of the family that is present in that genotype.

**Figure 3.**
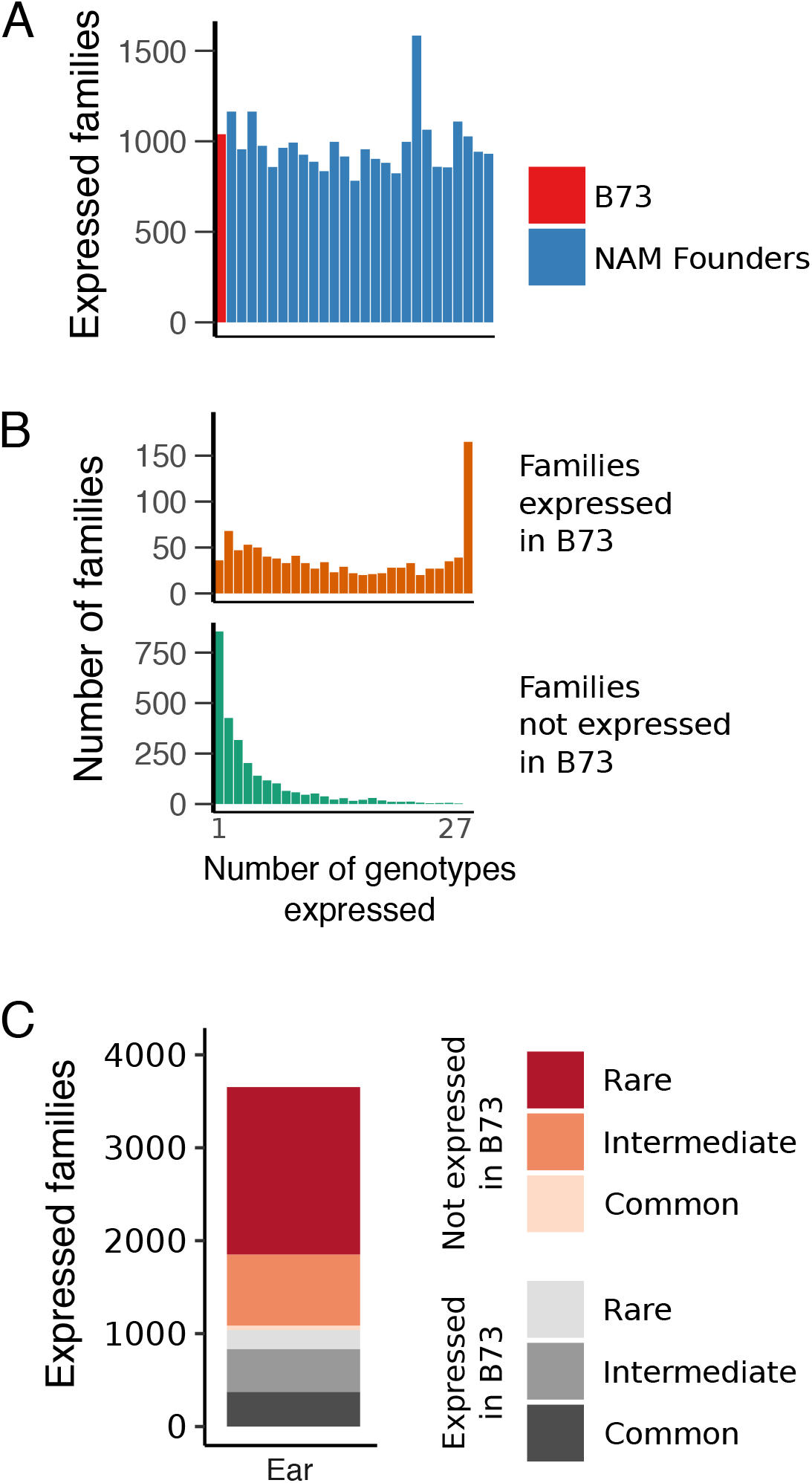

### TE expression in recombinant inbred lines

In order to perform a more detailed analysis of the role of TE family size variation and polymorphisms in the expression variation among genotypes we performed additional comparisons of TE transcript abundance in maize genotypes B73 and Mo17 as well as a set of 105 recombinant inbred lines (RILs) derived from them (Li *et al.* 2013). A previous comparison of TE presence/absence variation between these genotypes identified both shared and unique elements and revealed that ~40% of TEs in each genotype were absent in the other, totalling > 240,000 elements that were not shared between genotypes (Anderson *et al.* 2019). Transcript abundance of TE families (based on alignments of reads to the B73 genome/annotations) was assessed in shoot apex tissue for the two parents in addition to the RILs, and differentially expressed TE families between B73 and Mo17 were determined using DEseq2 (Fold-change >= 2, FDR < 0.05). This identified 278 TE families expressed higher in B73 and 239 families expressed higher in Mo17, including 95 and 98 families expressed only in B73 or Mo17, respectively. For all DE families, we assessed the relationship between the log2FC and the change in copy number between B73 and Mo17. This revealed minimal correlation between expression differences and the variation in the number of elements in the family across genotypes (Figure S5A). One potential explanation for this observation is that family-level expression is often determined by a single expressed member rather than equal contribution from all members of the family. To assess this, we looked at the distribution of RILs with expression for multi-member TE families expressed specifically in B73. This revealed that the majority of these families have expression in approximately half of the RILs, consistent with a single transcribed element segregating in the population (Figure S5B).

To identify the range of patterns for expression segregation in RILs, all TE families differentially expressed between B73 and Mo17 were assessed. Hartigan’s dip test index (Hartigan and Hartigan 1984) was used to determine the probability that expression values in the RILs exhibits a unimodal distribution (see methods for details). In this analysis, a single expressed locus that is segregating among the lines is expected to have a bimodal distribution, whereas families with several expressed members contributing quantitatively to expression are expected to show a unimodal distribution when segregating in the RILs. Across all DE families, ~20% have a unimodal distribution among RILs, and the proportion increases with larger family sizes in B73 (Figure S5C). An increased proportion of unimodally-distributed expression in the RILs was also found for families up-regulated in Mo17 and those with more members in Mo17 than B73.

Several example families were assessed in detail. A strong bimodal distribution among RILs is seen for family RLG05892, which is present as a single copy in B73 and absent from Mo17 (Figure 4 A-B). As expected, the vast majority of reads mapping to this family map uniquely to the single element (Figure 4 C-D). Interestingly, a bimodal distribution and expression of a single element is also seen for family RLG11255, which has a single, shared member in B73 and Mo17 (Figure 4). In total, there were 89 families differentially expressed between parents that have entirely shared elements in the two genomes, suggesting a role for epigenetic influences acting on shared TEs. In contrast, some families exhibit unimodal expression patterns as exemplified by RLG01150, which has 4 members in B73 and 12 members in Mo17. Here, unique reads can be assigned to 3 of the members in B73, though the presence of additional copies in Mo17 and the quantitative variation in expression suggests that more family members are likely mapping to the limited loci in B73.

**Figure 4.**
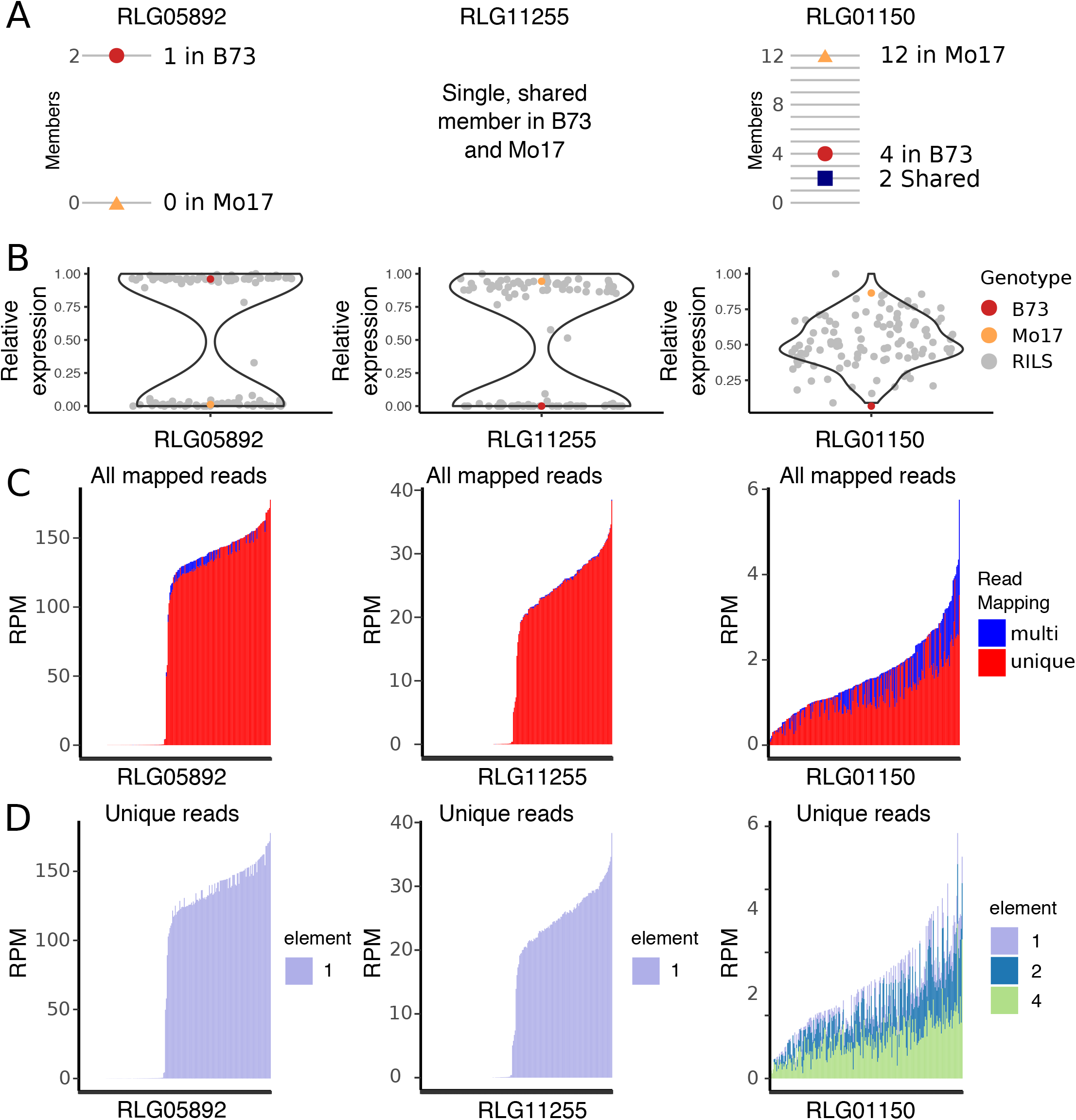

### TE expression in hybrids

In maize and other species, hybrids between distantly related lines can create heterosis defined by increased vigor relative to the parents. While the molecular cause of heterosis remains elusive, it has been suggested that combining substantially different complements of TEs could contribute to heterosis (Freeling *et al.* 2012). Given the potential for novel complements of TEs and sRNAs in the two parents to lead to novel regulation of TEs in the F1, we wanted to assess how TE expression changes in hybrids compared to inbred parents. Per-family TE expression was evaluated for trios containing B73, Mo17, and the F1 hybrid for 23 tissues of maize. For families with differential expression between B73 and Mo17, the deviation from additive expression was calculated (hybrid/mid-parent expression) and plotted for four tissues (Figure 5). The distribution of values for TE families centers around 0, consistent with the expression pattern seen for genes in these samples, suggesting largely additive expression patterns for TE families that are differentially expressed in B73 and Mo17.

**Figure 5.**
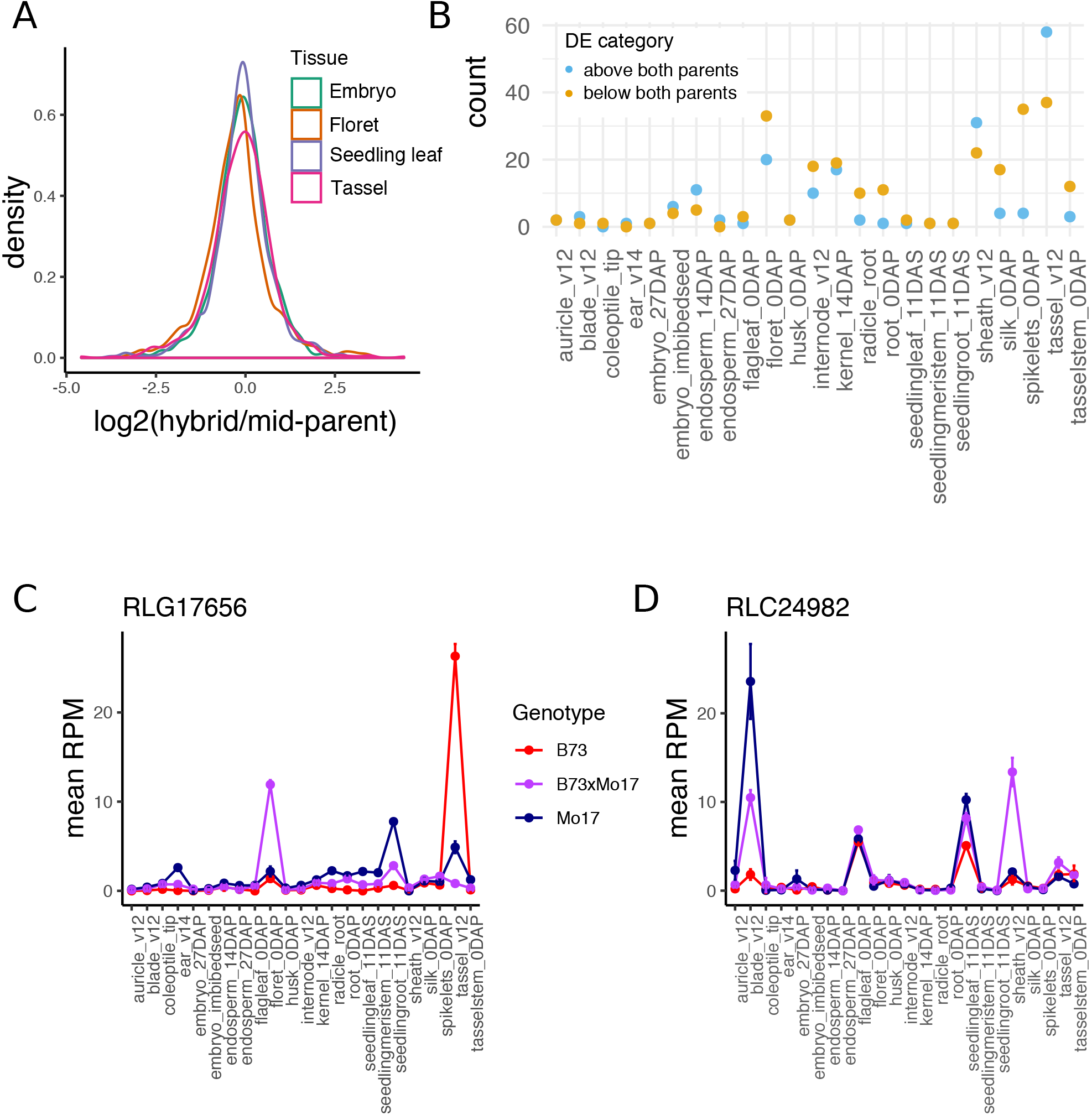

To identify TE families with significantly higher or lower expression than both parents (non-additive families), differential expression analysis was performed for each pairwise contrast using DEseq2 (Fold-change >= 2, FDR < 0.05). The number of non-additive families was determined for each tissue (Figure 5B), revealing that while many tissues have no examples of non-additive TE expression, some tissues, particularly inflorescence tissues, have a small number of families (< 5% of total expressed) showing non-additive expression. Closer inspection reveals that in many cases, non-additive expression is restricted to a single or a small number of tissues, with the TE family expressed higher in B73 or Mo17 in other tissues (Figure 5 C-D). This pattern of unstable non-additive expression across tissues is consistent with the observations for gene expression in these samples (Zhou *et al.* 2019). The breakdown of TE orders for families with non-additive expression is similar to the breakdown for all expressed families (83% LTR, 7% TIR, and 8% Helitron).

## Discussion

In this study, we assessed per-family TE expression in > 800 RNA-seq libraries representing tissue and genotypic diversity in maize. Although only a small proportion of TEs are ever expressed and total TE expression constituted only a small proportion of the transcriptome, TE families that were expressed are highly dynamic across both tissues and genotypes. In contrast to genes which exhibit constitutive or tissue-specific expression at similar rates, expressed TE families were almost exclusively tissue-specific. The dynamic nature of TE expression extends when assessing different genotypes, where the majority of expressed TE families are expressed in fewer than 20% of assessed genotypes. TE insertions are variable across different genotypes, and some of the variation in expression across genotypes can be attributed to variation in copy number and to the segregation of expressed families in populations. Finally, we found that while the majority of TE expression in hybrids is within the range of parental expression, a small number of families have expression significantly above or below both parents in some tissues, particularly tissues associated with male reproduction.

TEs contribute repetitive sequences to the genome resulting from both repetitive ends of elements (for example long terminal repeats for LTR elements) and proliferation into families with repetitive sequence. Due to these repetitive sequences, RNA-seq reads cannot always uniquely map to the genome, resulting in an underestimation of the total TE contribution to the transcriptome. We have developed a method that circumvents this problem by assessing TE expression per family rather than per element. While there are multiple ways to consider TE expression without throwing away multi-mapped reads (Slotkin 2018), we chose to assess family-level expression in order to best capture cases where a TE family contains a regulated element, resulting in coordinated expression of multiple family members.

In this work we have assessed TE expression by mapping reads to structural annotations of TEs. It is worth noting that the presence of transcripts that align to TE sequences may not represent expression of functional TE products. A subset of the expression we have observed likely does provide expression of the full retrotransposons or coding regions of TIR elements. However, in other cases the expression may be the result of a cryptic promoter within a TE and visualization for some of the TEs reveals that transcripts are only observed for a small region of the element rather than the full element. It is difficult to fully separate these types of expression for all TEs using short-read data. The application of long-reads for sequencing of full transcripts will be useful in revealing the contribution of different types of transcripts to the complement of TE expression. Even though some of the TE expression we have observed may not relate to production of functional transposon products it can be useful in providing information on the potential for TE regulatory influences in the genome. In some cases, regulatory elements within TEs have been shown to influence expression of nearby genes, either through acting as enhancers or creating merged transcripts initiating within the TE. Indeed, certain TE families in maize are associated with genes that show up-regulated expression in response to abiotic stress (Makarevitch *et al.* 2015) and this may reflect a more general mechanism through which regulatory elements are moved around the genome. By documenting the dynamic expression patterns of different TE families we can potentially gain insight into the regulation of the TE promoters and some of these regulatory influences may also affect the expression of nearby promoters.

While prior studies have reported increased TE expression in the male germ line, we find that the relative proportion of TE reads to gene reads in pollen and anthers is similar to other tissues. What is noteworthy about TEs in the male germline is that there are a large number of TE families with expression specifically in pollen and pollen-containing tissues (anther and floret). In *Arabidopsis*, TE de-repression in pollen predominantly occurs in the vegetative cell: a terminal tissue with close contact to the germ line (Slotkin *et al.* 2009). The other terminal but germline-adjacent tissue is the endosperm. There, TE reads do contribute proportionally more to the transcriptome than other tissues, however this is primarily due to high accumulation of a single TE family rather than global up-regulation of all TEs. Similarly to pollen, a number of TE families are expressed specifically in the endosperm and not in other vegative tissues or the adjacent embryo. It is interesting to note that there is not a group of TE families that were up-regulated in both male and female germlines, suggesting the possibility for a division of labor in germline TE silencing. A recent study in maize identified several TE families that were mobile only in the paternal germline in specific maize inbred genotypes (Dooner *et al.* 2019). However, no new TE insertions were identified in crosses where B73 was the pollen donor, so we were unable to assess how steady-state transcripts relates to known mobile elements.

The analysis of TE expression in multiple inbred genotypes of maize reveals substantial variation. There are many examples of TE families that show expression in some inbreds but not others. These differences are not strongly associated with the TE copy number of the family. Instead our analyses suggest that expression differences among genotypes reflect either differences in the presence/absence of a specific family member or changes in regulation of a shared TE. While there are examples of TE families in which many members are expressed there are also many examples in which expression of a TE family is due to expression solely from a single member of the family. In many cases where a TE family is differentially expressed between genotypes, we find that the member of the family that is expressed in B73 is missing in the genotypes without expression. However, in some cases we find that this element is present in both genomes but shows a difference in regulation. These findings suggest that both TE polymorphisms and regulatory variation, likely including epigenetic variation, can contribute to the observed differences in TE expression between genotypes.

Luxuriant TE expression in hybrids has been proposed as a source of hybrid vigor (Freeling *et al.* 2012). However, we find that the vast majority of expressed TE families do not show highly non-additive expression patterns in hybrids, and in fact there were more cases of families that were expressed much lower than both parents in hybrids than are expressed higher. While it is possible that TE mis-regulation is contributing to some of the phenotypic differences in hybrids, this is unlikely to result from global changes in TE expression. However, particular TEs or families may still contribute to the unique characteristics of hybrids.

## Acknowledgements

This work was funded by grants from USDA-NIFA2016-67013-24747 (S.N.A., C.D.H., and N.M.S.), NSF GRFP (M.C.S), NSF IOS-1546899 (P.Z. and N.M.S.) and NSF IOS-1238014 (J.R.I.). The Minnesota Supercomputing Institute (MSI) at the University of Minnesota provided computational resources that contributed to this research.

---

**Figure S1.**
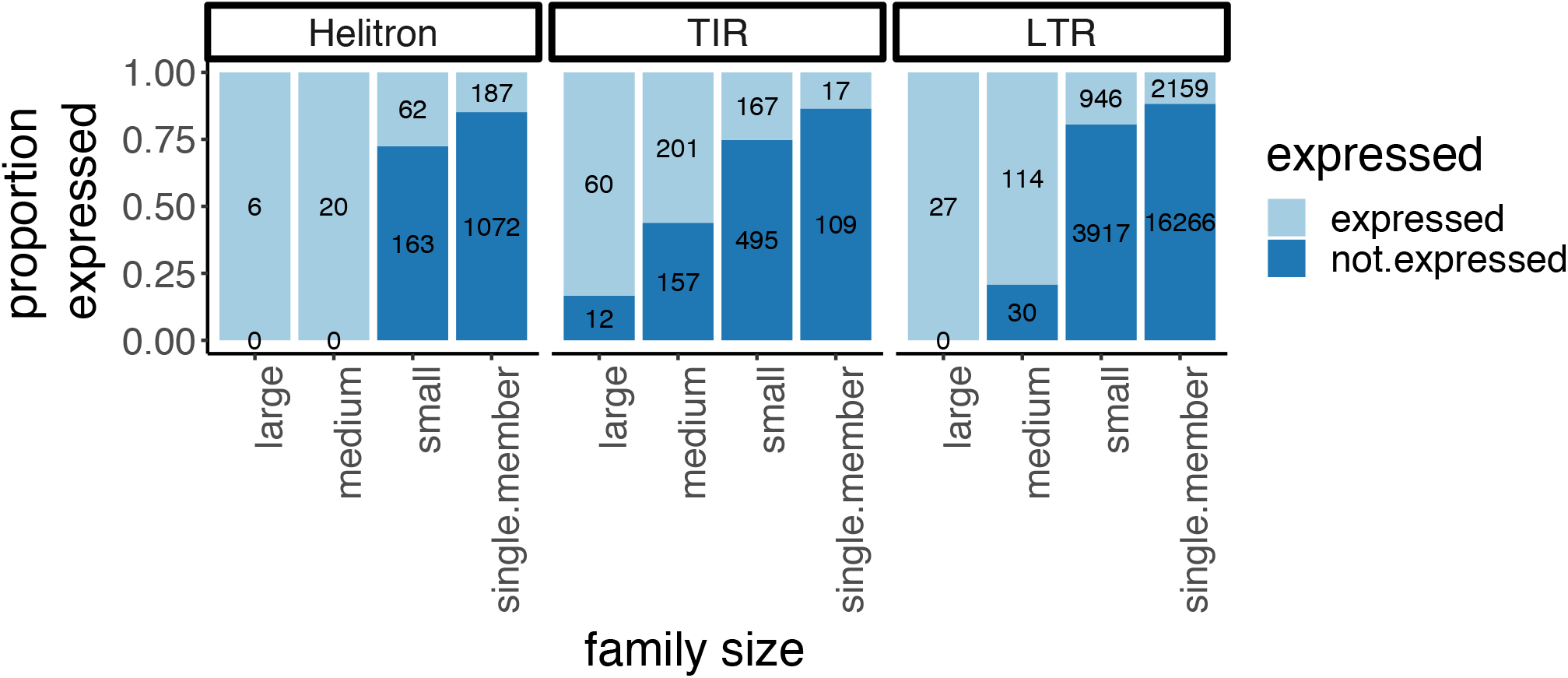

**Figure S2.**
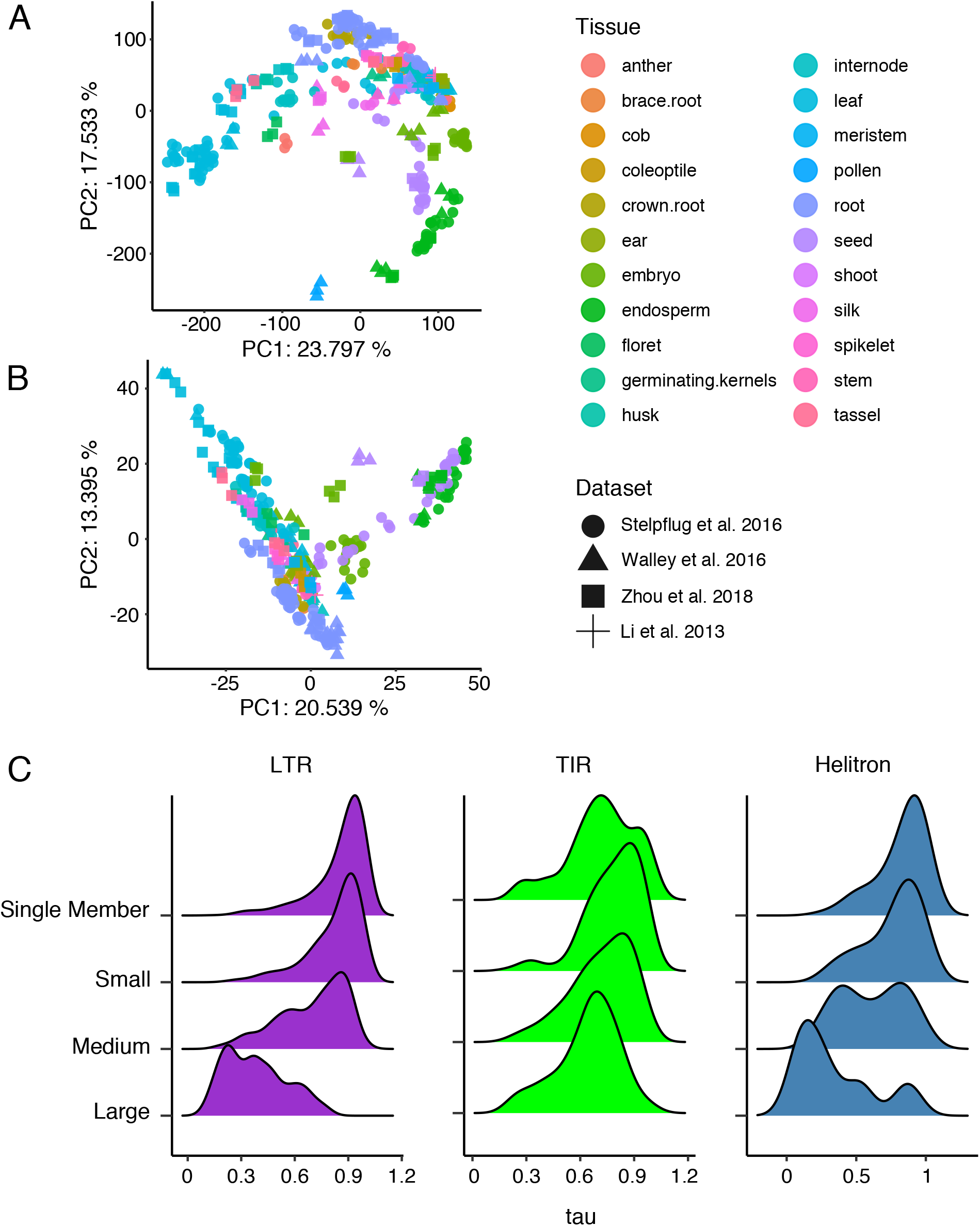

**Figure S3.**
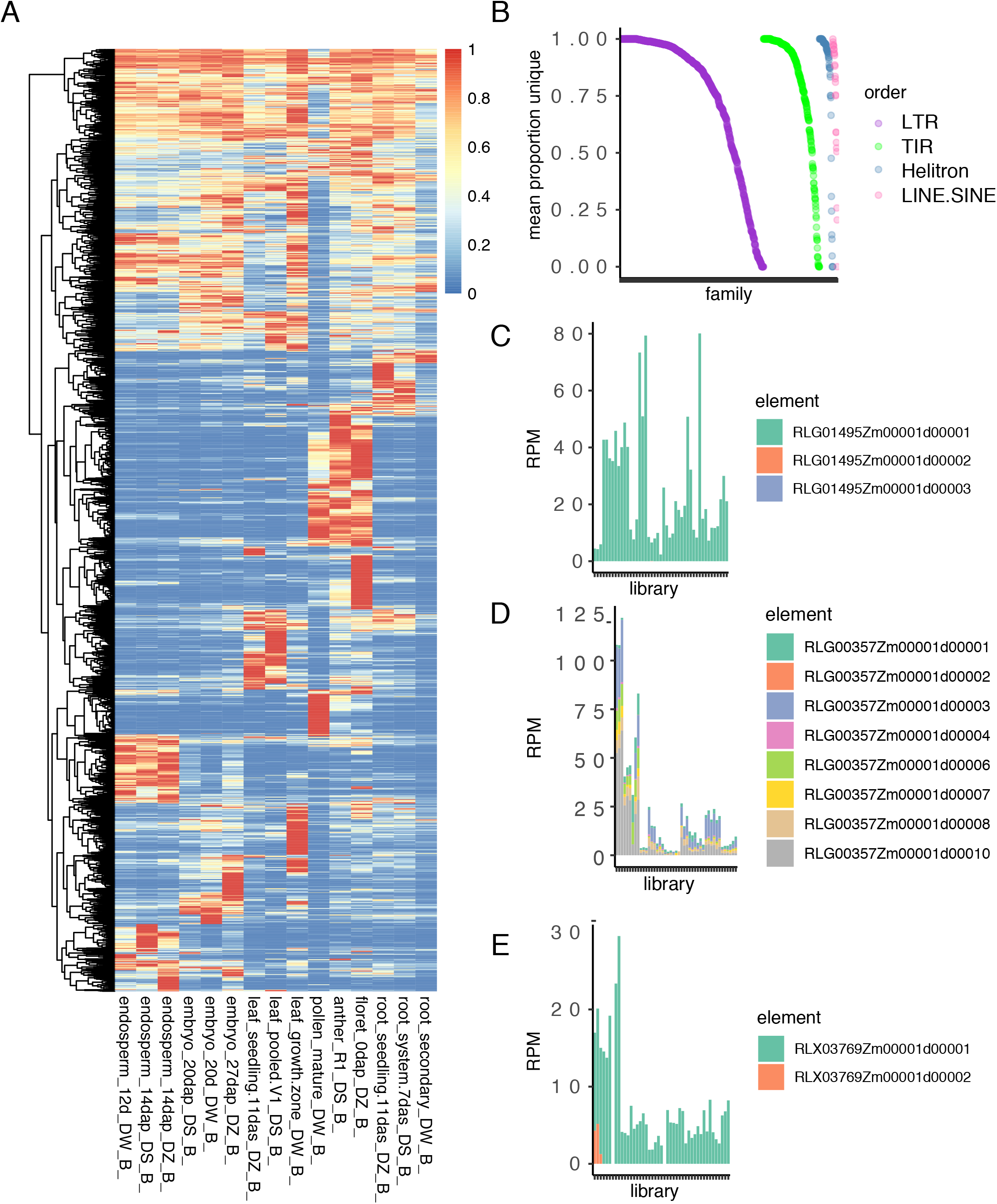

**Figure S4.**
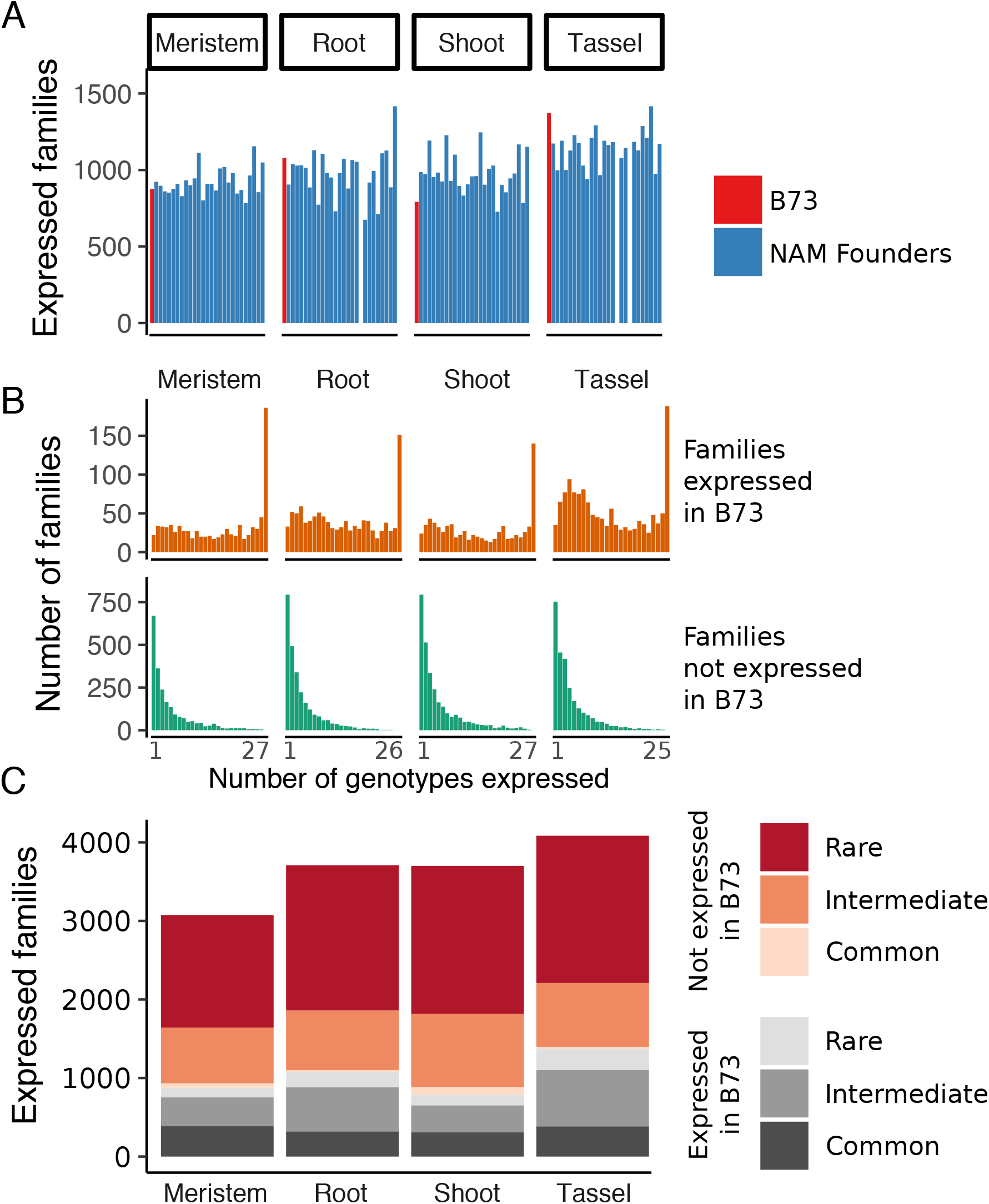

**Figure S5.**
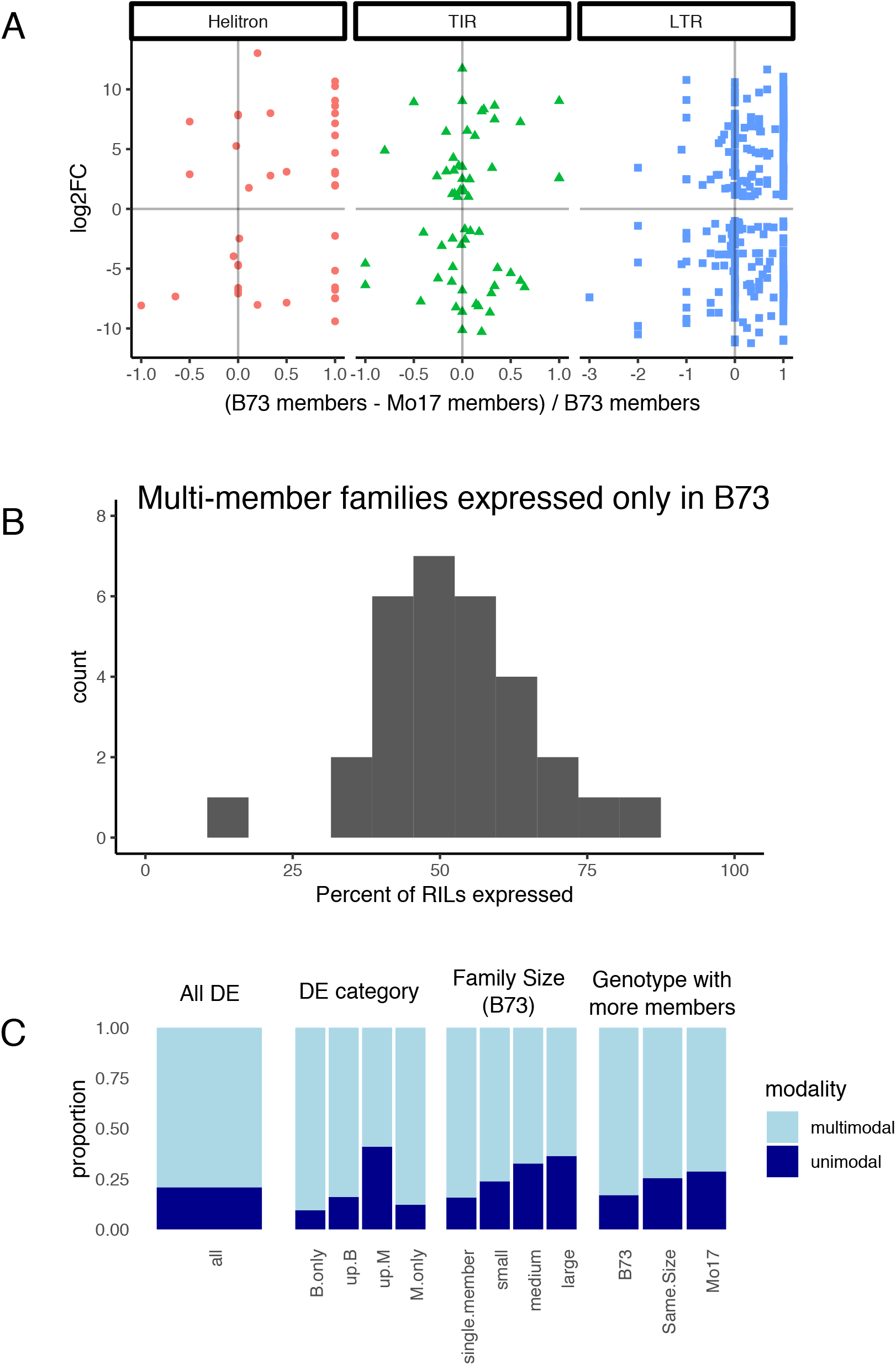

